# Deterministic influences exceed dispersal effects on hydrologically-connected microbiomes

**DOI:** 10.1101/088518

**Authors:** Emily B. Graham, Alex R. Crump, Charles T. Resch, Sarah Fansler, Evan Arntzen, David W. Kennedy, Jim K. Fredrickson, James C. Stegen

## Abstract

Subsurface groundwater-surface water mixing zones (hyporheic zones) have enhanced biogeochemical activity, but assembly processes governing subsurface microbiomes remain a critical uncertainty in understanding hyporheic biogeochemistry. To address this obstacle, we investigated (a) biogeographical patterns in attached and waterborne microbiomes across three hydrologically-connected, physicochemically-distinct zones (inland hyporheic, nearshore hyporheic, and river); (b) assembly processes that generated these patterns; (c) groups of organisms that corresponded to deterministic changes in the environment; and (d) correlations between these groups and hyporheic metabolism. All microbiomes remained dissimilar through time, but consistent presence of similar taxa suggested dispersal and/or common selective pressures among zones. Further, we demonstrated a pronounced impact of deterministic assembly in all microbiomes as well as seasonal shifts from heterotrophic to autotrophic microorganisms associated with increases in groundwater discharge. The abundance of one statistical cluster of organisms increased with active biomass and respiration, revealing organisms that may strongly influence hyporheic biogeochemistry. Based on our results, we propose a conceptualization of hyporheic zone metabolism in which increased organic carbon concentrations during surface water intrusion support heterotrophy, which succumbs to autotrophy under groundwater discharge. These results provide new opportunities to enhance microbially-explicit ecosystem models describing hyporheic zone biogeochemistry and its influence over riverine ecosystem function.

**Originality-Significance Statement:** Subsurface zones of groundwater and surface water mixing (hyporheic zones) are hotspots of biogeochemical activity and strongly influence carbon, nutrient and contaminant dynamics within riverine ecosystems. Hyporheic zone microbiomes are responsible for up to 95% of riverine ecosystem respiration, yet the ecology of these microbiomes remains poorly understood. While significant progress is being made in the development of microbially-explicit ecosystem models, poor understanding of hyporheic zone microbial ecology impedes development of such models in this critical zone. To fill the knowledge gap, we present a comprehensive analysis of biogeographical patterns in hyporheic microbiomes as well as the ecological processes that govern their composition and function through space and time. Despite pronounced hydrologic connectivity throughout the hyporheic zone—and thus a strong potential for dispersal—we find that ecological selection deterministically governs microbiome composition within local environments, and we identify specific groups of organisms that correspond to seasonal changes in hydrology. Based on our results, we propose a conceptual model for hyporheic zone metabolism in which comparatively high-organic C conditions during surface water intrusion into the hyporheic zone support heterotrophic metabolisms that succumb to autotrophy during time periods of groundwater discharge. These results provide new opportunities to develop microbially-explicit ecosystem models that incorporate the hyporheic zone and its influence over riverine ecosystem function.

## Introduction

Groundwater-surface water mixing zones (i.e., hyporheic zones) are characterized by temporally and spatially dynamic, redox-active environments with elevated rates of biogeochemical cycling (Hedin et al., 1998; McClain et al., 2003; Hancock et al., 2005; Boulton et al., 2010). Yet, hyporheic microbiomes have rarely been examined (Robertson and Wood, 2010; Marmonier et al., 2012; Gonzalez and Bell, 2013), constituting a key uncertainty in understanding the biogeochemistry of these dynamic zones (Robertson and Wood, 2010; Marmonier et al., 2012; Gonzalez and Bell, 2013). In particular, microbial community assembly processes drive microbiome composition and are thought to be imperative in coupling microbiomes with ecosystem function (Ferrenberg et al., 2013; Nemergut et al., 2013; Graham et al., 2016a; Graham et al., 2016b). Deterministic assembly (*e.g.*, selection) may either sort species such that the microbiome is optimized for a given environment or exclude taxa that may otherwise enhance ecosystem function (Knelman and Nemergut, 2014; Graham et al., 2016a), while stochasticity (*e.g.*, dispersal, drift) can regulate microbiome composition and function via mechanisms such as such as priority effects (Fukami, 2004; Fukami et al., 2010) and dispersal limitation (Lindström and Langenheder, 2012; Adams et al., 2013; Cline and Zak, 2014). Knowledge on the balance of assembly processes in determining microbiome composition and function in response to hydrologic exchange in the hyporheic zone is critical to understanding river corridor function, as hyporheic zones account for up to 95% of riverine ecosystem respiration (Naegeli and Uehlinger, 1997).

Determinism has been widely shown to impact environmental microbiomes (Fierer and Jackson, 2006; Lauber et al., 2009), while stochastic processes can strongly influence microbial community composition when environmental heterogeneity and/or species sorting is low (Stegen et al., 2012; Wang et al., 2013; Woodcock et al., 2013; Battin et al., 2016). Both of these processes operate in concert to structure microbiomes, and recent work has focused on deciphering the balance of deterministic vs. stochastic assembly processes through space and time (Dini-Andreote et al., 2015; Gianuca et al., 2016; Graham et al., 2016a). Further, assembly processes function at an organismal level; determinism selects for traits contained within certain groups of organisms and dispersal disproportionately affects organisms expressing traits for motility (Martiny et al., 2006; Martiny et al., 2013). Understanding microbiome assembly at both the community and sub-community levels is therefore essential to fully comprehending microbiome assembly and subsequent ecosystem functioning. Nonetheless, influences of assembly processes on specific groups of microorganisms are poorly understood, and linking a process-based understanding of variation in microbial physiologies to the metabolism of carbon and nutrients remains a vital unknown in ecosystem science.

The role of community assembly processes in determining microbiome composition and function may be particularly important within hyporheic zones that experience frequent groundwater-surface water mixing (hereafter, hydrologic mixing) and harbor a variety of complementary resources (Boulton et al., 2010; Stegen et al., 2016). For instance, dispersal may be paramount to structuring hyporheic microbiomes, as hydrologic mixing induces physical transport of microbial cells across physicochemically-distinct environments. A heightened importance of stochasticity has been hypothesized to generate a disconnect between microbiome composition and biogeochemical function (Nemergut et al., 2013; Graham et al., 2016b), and such a scenario would suggest microbiome composition is critical to accurately predicting biogeochemical function. Although most biogeochemical activity occurs within hyporheic sediments (Boulton et al., 1998; Hancock et al., 2005; Boulton et al., 2010), waterborne hyporheic microbiomes serve as dispersal vectors among attached microbiomes. Both types of microbiomes therefore convey mechanistic information regarding ecological dynamics that influence hyporheic function. Finally, strong local selective filters may negate dispersal effects by limiting successful immigration into physicochemically-distinct environments. In such a case, gradients in physicochemistry both determines microbiomes and drives hyporheic biogeochemical cycling (Graham et al., 2014; Graham et al., 2016b).

Our research has four primary objectives: to determine (a) biogeographical patterns in hyporheic microbiomes across three physicochemically-distinct zones and to infer associated trends in microbial physiologies; (b) assembly processes that generate these patterns through space and time; (c) specific groups of organisms whose relative abundances are deterministically driven by changes in the environment; and (d) how each identified group corresponds to hyporheic biogeochemical function. We hypothesize that (a) physicochemical differences across zones and microbiomes types (i.e., attached and waterborne) generate differences in microbiome composition and physiology; (b) hydrologic connectivity facilitates microbial dispersal across hyporheic zones despite steep physicochemical gradients; (c) the relative abundance of heterotrophic vs. autotrophic organisms corresponds to changes in organic carbon concentration associated with hydrologic mixing conditions; and (d) increases in the relative abundance of heterotrophic taxa corresponds with enhanced hyporheic zone biogeochemical activity.

## Results

### Hydrology and Physicochemical Conditions

The Hanford Reach of the Columbia River exhibits frequent hydrologic mixing and steep physicochemical gradients, presenting a prime opportunity for investigating the impacts of assembly processes on environmental microbiomes. We sampled distinct but hydrologically-connected zones that lie in close proximity (<250m) – the inland groundwater aquifer, the nearshore subsurface environment, and the Columbia River surface water – across seasonally variable hydrologic mixing. We observed variation in temporal dynamics of hydrologic mixing and physicochemistry among zones (Fig. 1A-C, Table S1, Fig. S1). Increases in chloride (Cl^-^) concentration in nearshore and inland zones at and beyond our July 22 sampling event indicated discharge of groundwater through the hyporheic zone (Graham et al., 2016a, Stegen et al. 2016, Fig. 1A), coincident with decreases in non-purgable organic carbon (NPOC) concentration in all three zones (Fig. 1B). Temperature followed a seasonal trend in nearshore and river zones but remained stable in the inland zone (Fig. 1C).

**Figure 1.**
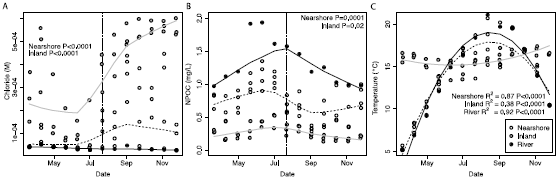
Changes in waterborne (A) Cl^-^ concentration, (B) NPOC concentration, and (C) temperature through time in each zone are presented in Figure 1. Lowess smoothers are plotted in panels (A) and (B) and quadratic regressions are plotted in (C) to aid in visualization. Data from nearshore, inland, and river zones are plotted with open, gray, and black circles, respectively. River stage dynamics are presented in Figure S1.

### Microbiome composition

River (PERMANOVA, R^2^ = 0.22, P = 0.001), nearshore attached (PERMANOVA, R^2^ = 0.44, P = 0.001), nearshore waterborne (PERMANOVA, R^2^ = 0.44, P = 0.001), and inland waterborne (PERMANOVA, R^2^ = 0.41, P = 0.001) microbiomes changed seasonally. Inland attached microbiomes (PERMANOVA, P = 0.29) were stable through time. Comparing all 5 data subsets to each other revealed distinct microbiomes (PERMANOVA R^2^ = 0.25, P = 0.001, Fig. S3), and within each zone the composition of waterborne and attached communities differed (PERMANOVA, nearshore R^2^ = 0.19, P = 0.001, inland R^2^ = 0.14, P = 0.001).

The ten most abundant phyla and classes in each data subset are presented in Table 1. Members of *Proteobacteria* were ubiquitous, and *Bacteroidetes* was highly abundant in all river and nearshore subsets but had lower abundance in inland zone habitats. The *Planctomycetes-Verrucomicrobia-Chlamydiae* (PVC) superphylum was prevalent, though the composition of organisms within this phylum varied among data subsets. *Chloracidobacteria* was widespread in attached microbiomes in both nearshore and inland zones but was not highly abundant in waterborne microbiomes. In the inland zone, classes *PBS-25* and *koll11* of the *OP3* candidate phylum were among the ten most abundant classes in the waterborne microbiome. Further, at the class level, the ammonia-oxidizing *Thaumarcheaota* was highly abundant in both inland waterborne and attached microbiomes, joined by a nitrite-oxidizer (*Nitrospira*) in attached microbiomes.

**Table 1.**
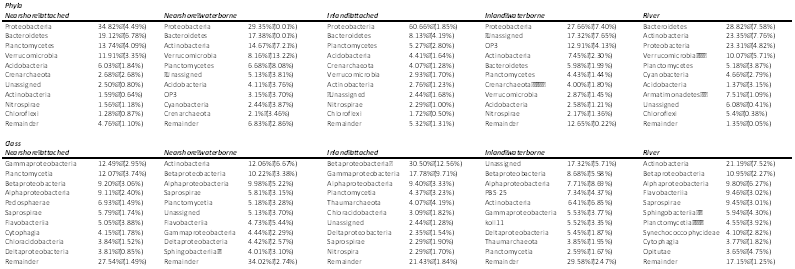
Phyla and classes with the highest relative abundance during our study period for each data subset. Mean relative abundance is presented with standard deviations listed in parentheses.

### Microbiome assembly

We used ecological modeling to infer the relative influences of deterministic selection vs. stochastic processes on microbiome composition—and futher parsed out the influences of high and low levels of dispersal within the stochastic component—in attached and waterborne communities. Deterministic assembly processes had more influence on microbiome composition than stochastic processes (Table 2). Below, we refer to homogenous selection (i.e., consistent selection pressures between locations or points in time), variable selection (i.e., divergent selection pressures between locations or points in time), dispersal limitation (i.e., inhibition of transport between microbiomes coupled with stochastic demographics), and homogenizing dispersal (i.e., heightened transport between microbiomes, leading to establishment) as per definitions outlined by Stegen et al. (2015). In attached microbiomes, both nearshore and inland, homogenous selection accounted for 82-100% of assembly processes, and for river microbiomes homogenous selection account for >95% of assembly. While waterborne microbiomes in nearshore and inland zones exhibited higher stochasticity than their attached counterparts, selection still accounted for 59.8% and 35.7% of assembly, respectively (Table 2).

**Table 2.**
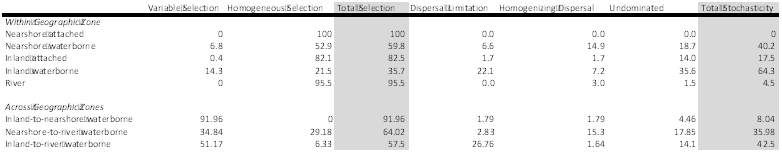
Assembly processes influencing microbiome composition as percentage of assembly mechanisms. Assembly processes are defined according to Stegen et al. (2015, see main text), with the “undominated” fraction being the residual fraction of comparions after accounting for all other assembly processes.

Because we hypothesized dispersal mechanisms to have the greatest impact on waterborne microbiomes compared across zones, we investigated assembly processes influencing across-zone comparisons within waterborne microbiomes only. Selection comprised 57-92% of across-zone assembly processes, with dispersal being of secondary importance. Variable selection was responsible for >90% of dissimilarity in microbiomes between nearshore and inland zones, while nearshore-to-river comparisons showed an even balance of variable (34.84%) and homogenous (29.18%) selection. The influence of spatial processes was most evident as dispersal limitation (26.76%) in inland-to-river comparisons and as homogenizing dispersal (15.30%) in nearshore-to-river comparisons.

### Spatiotemporal co-occurrence networks, environmental correlations, and keystone taxa

Finally, we used co-occurrence networks to infer relationships between deterministic assembly and specific groups (i.e., statistical clusters) of organisms, as well as to identify taxa vital in determining the composition of each cluster of organisms (i.e., keystone taxa). We also investigated relationships between each cluster of organisms and changes in hyporheic biogeochemical functioning by correlating the abundance of each cluster with active biomass (ATP) and aerobic respiration (resasurin assay, referred to below as ‘Raz’).

Properties of full networks at the family level are listed in Table S2, and the composition of clusters are listed in Table S3. In the nearshore zone, attached cluster 1 was negatively correlated with time, Cl-, and temperature (Fig. 2A-D); waterborne cluster 9 was positively correlated with time, Cl-, and temperature and negatively correlated with NPOC (Fig. 2E-H). In the inland zone, clusters of attached organisms did not display consistent trends with time or physicochemistry. Cluster 3 (Fig. 2I-L) and cluster 5 (Fig. 2M-P) of inland waterborne microbiomes exhibited contrasting relationships with time (positive/negative), Cl^-^ (positive/negative), NPOC (negative/positive), and temperature (positive/negative). River waterborne cluster 2 was negatively correlated with time and positively correlated with Cl^-^, with no evident relationships to NPOC or temperature (Fig. 2Q-T). Only one cluster-nearshore attached cluster 1-exhibited correlations with ATP and Raz (Fig. 3).

**Figure 2.**
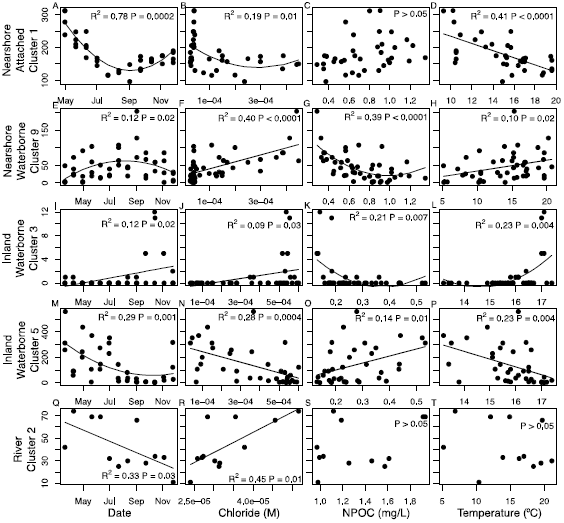
Regressions between the abundance of selected clusters and day of year (column 1), Cl^-^ concentration (column 2), non-purgable organic carbon (NPOC) (column 3), and temperature (column 4). Nearshore attached cluster 1 is presented in row 1 (panels A-D), nearshore waterborne cluster 9 is in row 2 (E-H), inland waterborne cluster 3 is in row 3 (I-L), inland waterborne cluster 5 is in row 4 (M-P), and river cluster 2 is in row 5 (Q-T).

**Figure 3.**
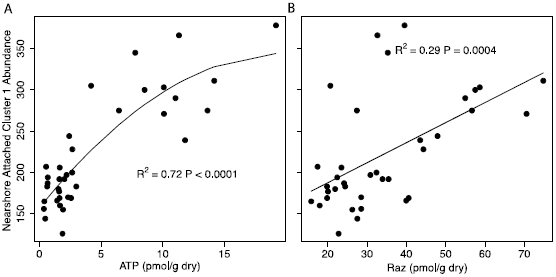
The abundance of nearshore attached cluster 1 was positively associated with (A) active microbial biomass and (B) aerobic respiration. Class-level analyses are presented here; family level-analyses are presented in Figure S2.

Nearshore attached cluster 1 contained keystone taxa belonging to *Verrucomicrobia* and *Thaumarcheaota*, as well as families of *Gammaproteobacteria*, *Alphaproteobacteria*, and *Chloracidobacteria* with a secondary importance (Fig. 4A, Table S3). Nearshore waterborne cluster 9 contained two keystone families—one family of unassigned organisms and one belonging to the candidate phylum *OP3* (Fig. 4B). No keystone taxa were identified in inland waterborne cluster 5 (Fig. 4C), but inland waterborne cluster 3 contained two keystone families belonging to *Chloracidobacteria* and *Chloroflexi* as well as organisms with secondary importance (Fig. 4D). No keystone taxa were identified in river cluster 2 (Fig. 4E).

**Figure 4.**
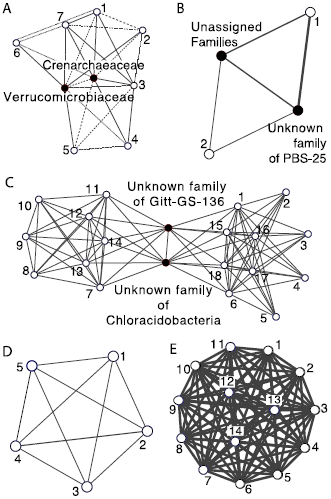
Network topology of clusters of organisms that correlated with time, Cl^-^, NPOC, temperature, ATP, and/or Raz (see Figure 3) are presented in (A) nearshore attached cluster 1, (B) nearshore waterborne cluster 9, (D) inland waterborne cluster 3, (E) inland waterborne cluster 5, and (E) river cluster 2. Edge thickness denotes the strength of Spearman?s correlation between two nodes (ranging from 0.6 to 1.0), and keystone taxa are denoted as black nodes with labels. All other nodes are numbered according to Table S3, and BC values for all nodes are also listed in table S3. Solid edges represent positive correlations and dashed edges represent negative correlations.

## Discussion

Hydrologic mixing conditions changed through time and corresponded to shifts in aqueous physicochemistry and microbiome composition across all zones (Fig. 1, Fig. S1). In the following sub-sections, we parse (a) biogeographical patterns in microbiomes, (b) microbiome assembly processes, (c) network clusters linked with seasonal changes in physicochemistry, and (d) associations between network clusters and hyporheic metabolism.

### Microbiome biogeography

Despite hydrologic connectivity across zones, distinct microbiomes were maintained within each zone and within attached vs. waterborne environments (Febria et al., 2012; Niederdorfer et al., 2016), supporting our hypothesis that physicochemical differences would lead to distinct microbiomes.

Among differences in microbiomes, we observed higher abundances of *Chloracidobacteria* in attached vs. waterborne microbiomes within both nearshore and inland zones (Table 1). This suggests a potentially important role for *Chloracidobacteria* in microbial biofilms attached to sediment particles. Biofilm communities, which are common in hyporheic zones, consist of a mixture of eukaryotic and prokaryotic organisms embedded in a porous extracellular matrix and contain diverse organisms with complementary metabolisms (Battin et al., 2016). At our sampling scale, we capture the diversity of organisms in biofilms but were unable to resolve spatial organization that may be key to elucidating specific relationships among these organisms. Still, *Chloracidobacteria* were prevalent in attached microbiomes (*i.e.*, within biofilms), possibly indicating a role in biofilm stability and functioning. These organisms are thought to be photoheterotrophs (Bryant et al., 2007; Tank and Bryant, 2015) that can utilize non-visible near-infrared light as an energy source for growth (Behrendt et al., 2012). While these properties may confer advantages in subsurface sediment conditions with low levels of light, our environments experience no light, and the prevalence of *Chloracidobacteria* suggests possible alternative physiologies for these organisms or dormant lifestyles to allow persistence in non-optimal environments (Gasol et al., 1995; Lennon and Jones, 2011).

Further, inland communities (characterized by a low organic C, high ionic strength environment) also contained many unique taxa relative to river and nearshore microbiomes, reflective of a distinct selective envrionment (Table 1). In particular, we observed members of the candidate phylum *OP3* in high abundance in the inland waterborne microbiome. *OP3* has been shown to be abundant in other subsurface communities with high inorganic C concentrations (Emerson et al., 2015) and appears to thrive in low oxygen environments, potentially utilizing anaerobic metabolisms reliant on iron, manganese, and/or sulfur (Glöckner et al., 2010). Our inland zone was characterized by high levels of inorganic C but remained oxygenated year round (Table S1), and although anaerobic microniches can form in interstitial water, the relative abundance of this taxon (∼13% of microbiome, 2^nd^ most abundant phylum) suggests that *OP3* may be able to persist under a broader range of redox conditions than currently recognized. Further, nitrifying organisms *Thaumarcheaota* and *Nitrospira* were prevalent in the inland zone. These organisms are chemosynthetic autotrophs, not dependent on external organic material, and instead fix CO_2_ through autotrophic pathways involving NH_4_^+^ and NO_2_^-^ oxidation. A higher level of variable selection in the inland zone relative to other zones (Table 2) is consistent with niche diversification, and the unique physicochemical environment of the inland zone may allow for diversification of oligotrophic organisms belonging to *OP3*, *Thaumarcheaota*, and *Nitrospira*, while excluding heterotrophic organisms that are dominant in other zones.

Despite pronounced biogeographical differences in microbiomes, our results indicate some common selective features and/or dispersal routes across zones and habitats. We observed high abundances of *Proteobacteria* and members of the PVC superphylum in all data subsets, and *Bacteroidetes* was among the most abundant phyla in both waterborne and attached microbiomes in nearshore and river zones (Table 1).

The ability of *Proteobacteria* to assimilate low-energy substrates, such as sulfate, nitrate, and inorganic C—in addition to various organic C compounds—may contribute to their abilities to thrive across physicochemically-distinct aquatic environments (Cottrell and Kirchman, 2000; Brenner et al., 2005; Battin et al., 2016). For example, *Alphaproteobacteria* and *Betaproteobacteria* exhibit diverse heterotrophic metabolisms (Sato et al., 2009; Newton et al., 2011; Battin et al., 2016). *Gammaproteobacteria* can fix inorganic C and digest organic C (Nikrad et al., 2014; Dyksma et al., 2016), while *Deltaproteobacteria* utilize organic C as carbon sources and as electron donors (Lovley et al., 1998; Torres et al., 2010). Microbiomes with a diverse community of *Proteobacteria* may therefore be well-adapted to fluctuations in organic C availability, facilitating their persistence across all zones. Moreover, *Bacteriodetes* has been shown to be prevalent in both waterborne and biofilm microbiomes in other systems—preferentially degrading recalcitrant C compounds and recycling biomass (Cottrell and Kirchman, 2000; Martin et al., 2015; Bennke et al., 2016). The presence of *Bacteroidetes* in addition to *Proteobacteria* within nearshore and river microbiomes may therefore indicate niche complementarity within these higher-C environments, possibly coincident with spatial organization that is not captured in our measurements.

Additionally, members of the PVC superphylum were abundant in all data subsets and possess distinctive physiologies. Many of these organisms have a proteinaceous membrane conveying antibiotic resistance (Fuerst and Sagulenko, 2011; Speth et al., 2012) and/or metabolize C1 compounds such as methane (Dunfield et al., 2007; Fuerst and Sagulenko, 2011; Sharp et al., 2013). Although the biogeochemical implications of selection for these organisms are unexplored, their unique ecology, ability to consume methane, and abundance within our system merits future investigating into their role in C cycling in hyporheic environments.

### Microbiome assembly

Hydrodynamics can be a driver for both stochastic and deterministic assembly processes, as advection can physically transport microorganisms across geographic barriers as well as generate changes in physicochemical selective environments. However, biogeographical patterns in microbiomes suggest a limited influence of dispersal relative to selection in our system. We empirically assessed assembly processes structuring microbiomes using ecological null models and, contrary to our hypothesis, our results indicate that selective pressures overwhelmed dispersal processes in structuring microbiomes.

In particular, attached communities were nearly entirely assembled by homogeneous selective pressures and displayed network topologies indicating a highly-structured microbiome (i.e., high clustering coefficient and low heterogeneity, Table S3). While our measurements do not provide sufficient resolution to describe fine-scale spatial organization of biofilms, our measurement scale likely captures multiple levels of organization and niche partitioning in each sample, allowing for inferences regarding community-level interactions. Our results are consistent with a recent review that noted the prevalence of niche-based assembly processes over stochasticity in structuring stream-associated biofilm microbiomes (Battin et al., 2016). Battin et al. (2016) suggest micro-niches in biofilms select for specific organisms and repel poorly-adapted immigrating species. Indeed, substrate features may impose strong environmental filters on microbiomes—sediment geochemistry (Carson et al., 2007; Jorgensen et al., 2012) and matrix structure (Vos et al., 2013; Breulmann et al., 2014) can select for traits that enhance attachment on a particular substrate.

Likewise, river microbiomes were assembled via homogeneous selection but were also differentiated from nearshore and inland zone microbiomes via a small influence of dispersal limitation (Table 2). Network structure was also consistent with a highly-organized river microbiome (*i.e*., high clustering coefficient and low heterogeneity, Table S3). A relatively stable geochemical environment in the river may provide a steady selective environment despite seasonal changes in temperature (Table S1, Fig. 1C). In particular, NPOC concentration was approximately 2-5 times higher in river water than the nearshore and inland waterborne environments. Furthermore, organic C associated with surface water intrusion has been shown to be correlated with short-term increases in deterministic assembly processes and changes in microbiome composition in our nearshore environment (Stegen et al., 2016). Thus, consistently high NPOC concentration in the river may impose environmental filters for waterborne heterotrophic organisms within river microbiomes, resulting in homogeneous selection through time. Moreover, distinct river physicochemistry and physical filtering of microorganisms (Brunke, 1999; Hartwig and Borchardt, 2014), as compared to nearshore and inland zones (Table S1), may limit successful colonization of river-associated taxa within other zones.

Although deterministic assembly processes were prevalent forces in structuring microbiomes, nearshore and inland waterborne microbiomes showed a greater influence of stochasticity than other zones. Within the nearshore, significant fluxes of both groundwater and surface water may contribute to spatial assembly processes (e.g., dispersal). We observed the impact of both dispersal limitation and homogenizing dispersal on nearshore waterborne microbiomes, suggesting that this zone may experience periods of comparatively low and high rates of hydrologic transport (i.e., hydrologic mixing and interstial flow). Furthermore, sampling locations for inland waterborne microbiomes were distributed across a broader spatial extent than any other zone, perhaps facilitating our ability to detect dispersal limitation in these samples.

When we examined assembly processes differentiating waterborne microbiomes across zones, we anticipated more impact from spatial processes due to an increase in geographic scale. In contrast to this expectation, selection was responsible for more than 50% of community dissimilarity in all comparisons. While we have suggested physical inhibition of microbial dispersal as a possible mechanism generating microbiome dissimilarity across zones, our analyses indicate that an overarching influence of selection generally outweighed influences of spatial processes. Our results indicate that although spatial processes play a role in shaping microbiomes, local physicochemical environments often limit the ability of physically-transported microorganisms to outcompete local biota.

Indeed, nearshore-to-inland zones comparisons suggested that distinct selective pressures in each environment were the dominant cause (>90%) of differences among waterborne microbiomes. Likewise, spatial processes were overwhelmed by selection in nearshore-to-river comparisons. Differences among these microbiomes were due to relatively even proportions of homogenous and variable selection, with no evident seasonal trends in the balance between these processes. Our system is characterized by pronounced geomorphic heterogeneity in the subsurface environment that creates preferential flow paths for hydrologic mixing (Johnson et al., 2015). The presence of both homogeneous and variable selection may indicate spatial variation in surface water intrusion or local biogeochemical conditions across our sampling locations. Homogenizing dispersal also had a small but detectable influence in nearshore-to-river comparisons, supporting some hydrologic transport of microorganisms between these zones.

Lastly, assembly estimates between the inland and river zones provided the strongest evidence for spatial processes in our system (42.5% stochasticity). In particular, inland-to-river comparisons yielded the largest impact of dispersal limitation (26.8%), supporting a decay of community similarity across increasing spatial distances (Green and Bohannan, 2006). Dispersal limitation is, therefore, likely to play a more significant role in subsurface microbiome assembly at larger spatial scales.

### Network clusters, keystone taxa, and seasonal physicochemical change

Given pervasive deterministic assembly, we used network analysis to identify specific clusters of organisms whose relative abundances changed deterministically with changes in the environment within each zone. Co-occurrence networks fragment into clusters that are sensitive to variation in hydrological regimes (Widder et al., 2014; Febria et al., 2015) and have been used to identify keystone taxa (González et al., 2010; Vick-Majors et al., 2014; Banerjee et al., 2015; Banerjee et al., 2016). Our sampling scale aggregates multiple niches within each sample, allowing us to infer niches with similar vs. contrasting responses to environmental stimuli. In general, our results were consistent with our hypothesis that higher NPOC concentrations in surface water would shift microbiomes towards heterotophy. Specifically, we found that the prevelance of heterotrophic organisms increased with increased surface water contribution in each zone, and below we discuss microbial physiologies in each data subset that corresponded to seasonal changes in hydrology.

In attached microbiomes no relationships were present in the inland zone, but we found that nearshore attached cluster 1 was favored during early season conditions with pronounced surface water intrusion and low temperature (Fig. 2A-D). Clusters of organisms can denote both positive and negative co-occurrence patterns, signifying similar or opposite (respectively) ecological dynamics influencing taxa. For instance, organisms sharing a cluster may constitute an ecological niche, whereby organisms either co-occur in similar environments due to shared environmental requirements or be involved in beneficial interactions with each other (Shi et al., 2016). Alternatively, clusters may contain organisms that are favored under opposing environmental conditions or have negative interactions (e.g., competition, predation). Nearshore attached cluster 1 contained two anticorrelated groups of organisms—group 1 consisted of heterotrophic organisms (*Oxalobacteraceae*, *Comamonadaceae*, *Verrucomicrobiaceae*, and *Flavobacteriaceae*) that decreased through time, while organisms in group 2 were primarily autotrophic oligotrophs (*Crenarchaeaceae*, *Ectothiorhodospiraceae*, *Pelagibacteraceae*, and families of *Alphaproteobacteria*) that increased through time. Because of the abundance of organisms in group 1 compared to group 2 (4498 vs. 1987 total OTU counts), group 1 dictated correlations between this cluster and the environment, supporting an association between heterotrophic organisms and periods experiencing relatively high-organic C surface water intrusion. However, as the abundance of cluster 1 declined in concert with groundwater intrusion into the nearshore environment (Fig. 3A-D), organisms in group 2 increased, putatively outcompeting heterotrophic organisms in group 1 under more oligotrophic conditions. The co-association of heterotrophic and autotrophic organisms within a single cluster may indicate comparatively strong tradeoffs between these lifestyles and merits further investigation into niche dynamics and competitive interactions between these taxa. Further, we identified primary keystone taxa as *Verrucomicrobiaceae* and *Crenarchaeaceae*, possibly denoting the principal heterotrophic and autotrophic organisms in groups 1 and 2, respectively.

In waterborne microbiomes, we observed relationships between clusters of organisms in all zones with hydrologic mixing and physicochemistry. Nearshore waterborne cluster 9 displayed a seasonal association with groundwater discharging conditions and high temperature (Fig. 2E-H). Organisms were assigned to families highly abundant in the inland zone (*e.g*., *Thaumarchaeota*, candidate phylum *OP3*, Table 1, Table S3), and keystone taxa, in particular, belonged to *OP3*. Trends in the relative abundance of this cluster may therefore denote enhanced dispersal from the inland zone during groundwater discharge and/or a change in selection favoring these organisms corresponding to physicochemical shifts in nearshore environment. The inland waterborne microbiome also revealed two clusters related to environmental change. Cluster 3 was present in low abundance, and increased with higher groundwater contribution and temperature (Fig. 2I-L), and contained facultative anaerobic microorganisms (*Lentisphaeria*, *Gemmatimonadetes*, *Anaerolineae*, *Alphaproteobacteria*, *Gammaproteobacteria*, *Chloroflexi*, *Clostridia*, *Spirochaetes*) and methylotrophs (*Betaproteobacteria*, Table S3). Such organisms are largely distinct from organisms found in the nearshore and river environments, supporting the inference that physical and geochemical isolation drives differences between microbiomes. In contrast, cluster 5 was associated with surface water intrusion and low temperature (cluster 5, Fig. 2M-P). An influx of surface water-associated NPOC (Fig. 1B) should favor heterotrophic microorganisms, and cluster 5 contained methanotrophs (*Methylacidiphilae*, *Betaproteobacteria*) and diverse *Actinobacteria* and *Acidimicrobiia* involved in C cycling, possibly denoting a shift towards heterotrophy under comparatively high-organic C conditions. While no keystone taxa were found in cluster 5, *Chloracidobacteria* and a family of *Chloroflexi* were identified in cluster 3, meriting further investigation into their role in the hyporheic environment.

Finally, we identified one cluster of diverse river organisms that was positively associated with Cl^-^ concentration, which may indicate significant groundwater discharge into the river (Fig. 2Q-T, Table S3). In particular, *Cytophagia* was the most abundant organism in this cluster and was common in the nearshore and river zones, but all other families in this cluster were primarily found in the inland zone. These inland taxa were, however, in low abundance in the river and may therefore indicate a small contribution of the inland microbiome to river communities under discharging condtions. No keystone taxa were found in this cluster.

### Linkages between network clusters and hyporheic metabolism

Finally, we further investigated the involvement of each network cluster in microbial metabolism and found only one cluster—cluster 1 of nearshore attached microorganisms—that correlated with active biomass (ATP) and aerobic respiration (Raz). Positive correlations between cluster 1 and both ATP and Raz (Fig. 3) indicate that organisms within this cluster are either responsible for biomass growth and aerobic nutrient cycling or are facilitated by conditions under which these processes are promoted. While skewed towards heterotrophy, nearshore attached cluster 1 contained organisms with a broad range of metabolic capabilities, only partially supporting our hypothesis that increases in heterotrophic organisms would generate higher rates of biogeochemical activity. Complementary microbial resource use and niche diversification resulting from physical heterogeneity in sediments has been proposed to increase the rate and diversity of compounds metabolized by attached organisms in other systems (Singer et al., 2010; Battin et al., 2016). Moreover, spatial organization and excretion of extracellular enzymes that can be utilized across organisms in biofilms can allow for complementarity that enhances rates of metabolism (Loreau et al., 2001; Naeem et al., 2012). Nearshore attached cluster 1 may therefore constitute the foundation (or portion thereof) of this complementary resource structure.

### Conclusions

Our research generates a framework describing the roles of selection and dispersal in influencing microbal physiologies within hyporheic environments (Fig. 5A-B). We advance that spatiotemporal patterns in ecological selection impose strong environmental filters in hyporheic zones, limiting successful immigration of organisms across physicochemical gradients. For example, in Fig. 5A, hydrologic processes transport microorganisms from two communities to a third location, and the establishment of dispersing organisms in the third location is strongly impacted by local selective filters. Each community consists of largely distinct organisms, but contains a small proportion of similar organisms common to both dispersing communities, as evidenced by high levels of determistic assembly processes in our system but overlapping taxa between each zone. These selective filters also change with hydrologic mixing, inducing a shift from heterotrophic to autotrophic metabolisms with a change from surface water-dominated to groundwater-dominated hydrology (Fig 5B). This inference is supported by associations of nearshore attached and waterborne clusters as well as inland waterborne clusters with hydrologic mixing. All correlations between these clusters show a descrease in heterotrophs and an increase autotrophic organisms associated with increased groundwater contributions to physicochemistry.

**Figure 5.**
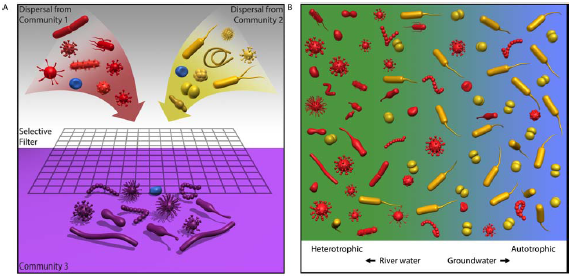
Conceptual diagrams depicting (A) influences of assembly processes on micriobiomes in our system and (B) microbiome shifts associated with hydrologic change in the nearshore hyporheic zone. In (A), hydrologic processes transport microorganisms from two communities to a third location. Each community consists of largely distinct organisms denoted by color (red vs. yellow), but contains a small proportion of similar organisms represented by the blue organism common to both dispersing communities. The establishment of dispersing organisms in the third location (purple zone) is strongly impacted by selective filters that limit successful immigration of organisms across environments. The local environment therefore selects for a unique microbiome (purple organisms) with the exception of a blue organism common to all three communities. Panel (B) depicts changes in microbiomes from heterotrophic to autotrophic metabolisms as hydrologic mixing conditions shift from surface water-dominated to groundwater-dominated. Mixing conditions are displayed as the background color, with green indicating surface water and blue indicating groundwater. Heterotrophic organisms are displayed in red, and autotrophic organisms are displayed in yellow.

Thus, despite largely deterministically-assembled microbiomes, a small proportion of common taxa across a continuum of groundwater-to surface water-dominated habitats indicate some influence of microbial dispersal on microbiome composition. We also identify organisms that change through seasonally dynamic hydrologic mixing conditions, providing insight into hyporheic microbial physiology. In particular, our research suggests a tradeoff between heterotrophic and autotrophic organisms under surface water-vs. groundwater-dominated hydrology. We therefore propose that comparatively high-organic C conditions during surface water intrusion into the hyporheic zone support heterotrophy, succumbing to autotrophy during time periods of groundwater discharge. Enhanced rates of activity associated with a nearshore attached cluster may also indicate a central biogeochemical role for taxa, and in particular keystone taxa (*Verrucomicrobiaceae and Crenarchaeaceae*), within this cluster. As a whole, our work shows that deterministic assembly may overwhelms dispersal processes in hyporheic environments despite pronounced hydrologic connectivity, and provides a process-based understanding of spatiotemporal patterns in microbiome composition and function in a zone of elevated biogeochemical cycling.

## Experimental Procedures

### Study Design

This study was conducted in three zones (inland, nearshore, river) within the Hanford 300A (approximately 46° 22’ 15.80”N, 119° 16’ 31.52”W) in eastern Washington State (Slater et al., 2010; Zachara et al., 2013; described in Graham et al., 2016a). Previous research has shown redox conditions as well as microbiome composition and activity to be variable across space and time in this system (Lin et al., 2012a; Lin et al., 2012b; Stegen et al., 2012; Graham et al., 2016a). sThe inland zone lies within 250m of the Columbia River and is characterized by an unconfined groundwater aquifer in the Hanford formation. It displays relatively stable temperatures (∼15°C) and elevated concentrations of anions and inorganic carbon relative to other zones (Table S1). In contrast, the Columbia River contains higher organic carbon and lower ion concentrations with seasonally variable temperatures. Surface water from the river and groundwater from the inland zone mix in the nearshore hyporheic zone. At high river stages, surface water intrudes into the inland zone; and at low river stages, groundwater discharges into the Columbia River. To monitor hydrologic mixing, we employed Cl^-^ as a conservative groundwater tracer per Stegen *et al*. (2016).

Detailed sampling and analytical methods are in the Supplemental Material. Briefly, attached (nearshore and inland only) and waterborne (all zones) microbiomes were obtained from deployed colonization substrate and waterborne samples, respectively. Colonization substrate was incubated *in situ* six weeks prior to removal. Samples were collected at three-week intervals from March through November 2014, with the first waterborne samples collected in March and the first attached samples collected in April after a six-week incubation period. Numbers of samples obtained at each timepoint are list in Table S4. Attached samples were obtained from fully screened stainless steel piezometers installed to 1.2m depth below the riverbed (nearshore, 5.25cm inside diameter (MAAS Midwest, Huntley, IL)) or from established groundwater wells (inland). Piezometers were installed across a zone of known groundwater-surface water exchange—and given the very significant cost of installing groundwater wells into the broader aquifer—we made use of existing groundwater wells that were as close as possible to the river shoreline. The need to use multiple pre-existing near-river wells resulted in a larger spatial extent covered by the wells than by the piezometers. While a larger inland spatial domain may have contributed to higher incidence of spatial assembly processes, this effect would only impact comparisons of within zone assembly processes (vs. across zone comparisons), and we note a dominance of selection within the inland zone similar to other zones (Table 2). Waterborne samples were obtained from galvanized piezometers located <1m from the piezometers used to sample attached microbiomes. For each inland location, a single well was used for both waterborne and attached samples. River water was sampled adjacent to the piezometers. DNA was extracted from each sample using the MoBio PowerSoil kit (MoBio Laboratories, Inc., Carlsbad, CA), and the 16S rRNA gene was sequenced on the Illumina MiSeq platform as described in the Supplemental Material. Physicochemical properties, active biomass (ATP), and aerobic respiration (Raz) were determined as per the Supplemental Material.

### Statistical analysis

All analyses were conducted in R software (http://cran.r-project.org/) unless otherwise noted. Relative abundances of major microbial taxa at phylum-and class-levels were calculated in each of five data subsets (nearshore attached, nearshore waterborne, inland attached, inland waterborne, river). An apparent breakpoint in hydrology at July 22 was assessed by comparing Cl^-^ and NPOC concentration pre-and post-July 22 with one-sided Mann Whitney U tests. Temporal changes in temperature in each subset were assessed with quadratic regressions. Microbiome dissimilarities were estimated as Bray-Curtis distance and analyzed through time and across data subsets with PERMANOVA in QIIME (Caporaso et al., 2010).

### Modeling assembly processes

We implemented null modeling methods developed by Stegen *et al*. (2013; 2015) to disentangle community assembly processes. The approach uses pairwise phylogenetic turnover between communities, calculated using the mean-nearest-taxon-distance (βMNTD) metric (Webb et al., 2008; Fine and Kembel, 2011), to infer the strength of selection. Communities were evaluated for significantly less turnover than expected (βNTI < -2, homogeneous selection) or more turnover than expected (βNTI > 2, variable selection) by comparing observed βMNTD values to the mean of a null distribution of βMNTD values—and normalizing by its standard deviation—to yield the beta-nearest taxon index (βNTI) (Stegen et al., 2012). Pairwise community comparisons that did not deviate from the null βMNTD distribution were evaluated for the influences of dispersal limitation and homogenizing dispersal by calculating the Raup-Crick metric extended to account for species relative abundances (RC_bray_), as per Stegen *et al*. (2013; 2015). Observed Bray-Curtis dissimilarities were compared to the null distribution to derive RC_bray_. RC_bray_ values that were > 0.95, > -0.95 and < 0.95, or < -0.95 were interpreted as indicating dispersal limitation, no dominant assembly process, or homogenizing dispersal, respectively. Significance levels for βNTI and RC_bray_ are respectively based on standard deviations—|βNTI| = 2 denotes two standard deviations from the mean of the null distribution—and alpha values—|RC_bray_| = 0.95 reflects significance at the 0.05 level. Inferences from both βNTI and RC_bray_ have previously been shown to be robust (Dini-Andreote et al., 2015; Stegen et al., 2015).

### Co-occurrence network analysis

Co-occurrence networks were constructed using Spearman’s correlation at four taxonomic levels – class, order, family, and OTU (singletons removed). To examine taxa with moderate to strong co-occurrence patterns, correlations with *rho* > +/-0.6 and FRD-corrected *P* < 0.01 were imported into Cytoscape v3.3 for visualization and calculation of network parameters. Networks were examined for deviations from randomness by comparing network parameters to random networks generated with Network Randomiser 1.1 in Cytoscape (Barabasi-Albert model with same number of nodes). Keystone taxa were identified using the Betweenness Centrality metric (BC > 0), whereby increasing BC values indicate greater contribution of nodes to network structure (González et al., 2010; Vick-Majors et al., 2014; Banerjee et al., 2016).

We identified clusters in Cytoscape using MCODE, using default parameters as per Banerjee *et al*. (2015; 2016). We then calculated Pearson’s momentum correlation between the relative abundance of each cluster and selected parameters (Cl^-^, NPOC, temperature, ATP, and Raz) to screen for clusters of interest (*P* < 0.05). Regression models (linear and quadratic, as appropriate) were then fit between cluster abundance (dependent variable) and the selected parameters (independent variables). Analyses were conducted at all taxonomic levels, but we present family-level correlations of microbiomes with time and physicochemistry due to the consistency of patterns. For correlations with microbiome activity and aerobic respiration, results with were similar at family-and class-levels (Fig. S2 and Table S5), and we present class-level results that exhibited better model fit.

## Acknowledgements

This research was supported by the US Department of Energy (DOE), Office of Biological and Environmental Research (BER), as part of Subsurface Biogeochemical Research Program’s Scientific Focus Area (SFA) at the Pacific Northwest National Laboratory (PNNL). PNNL is operated for DOE by Battelle under contract DE-AC06-76RLO 1830. A portion of the research was performed using Institutional Computing at PNNL.

